# Differential functions of upstream and intronic HOT regions shape *dlg-1* transcription and *C. elegans* development

**DOI:** 10.1101/2025.06.13.659077

**Authors:** Cristina Tocchini, Fanny Eggeler, Palmer Bassett, Susan E. Mango

## Abstract

High occupancy target (HOT) regions, genomic sites bound by many transcription factors, are found across eukaryotes, but their physiological roles remain unclear. We characterized two HOT regions in the *Caenorhabditis elegans dlg-1* locus containing transcription start sites: one upstream the coding sequence and one in the first intron. The upstream region is essential for proper transcription and proper animal development, while the intronic region modestly enhances transcription in epidermal cells. Our findings reveal the functions of HOT regions in animal development, expanding the repertoire of gene regulation strategies in *C. elegans*.

## Background

Precise regulation of gene expression is crucial for orchestrating complex developmental processes. Transcriptional control relies on a series of regulatory elements located within promoters or in distant genomic regions containing enhancers or repressors (1). In *Caenorhabditis elegans*, transcriptional regulation mainly relies on regulatory sequences within promoters, in the close proximity to a gene’s coding sequence (CDS). High occupancy target (HOT) regions are genomic sites bound by unusually large numbers of transcription factors (2,3). HOT regions have been identified across the *C. elegans* genome and may coincide with enhancer elements, promoters and transcription start sites (TSS) that commonly activate transcription ubiquitously (2,4,5). The contribution of HOT regions to transcriptional regulation in a physiological context remains poorly understood.

Here, we dissect the regulatory landscape of the *dlg-1/Discs large* locus during embryonic morphogenesis (6,7). Previously published genomic data revealed multiple candidate regulatory elements for the *dlg-1* gene (2,5,8). We focused our attention on two HOT regions marked by strong peaks of RNA Polymerase II (PolII) revealed by chromatin immuno-precipitation and sequencing (ChIP-seq), and containing TSS (5,9,10): one located upstream the CDS, and one in the first intron. Notably, the upstream HOT region is located more than four kilo-bases (Kb) away the CDS, and the one in the first intron contains enhancer sites, too. Long distances between TSS and CDS are likely allowed by widespread trans-splicing in *C. elegans*. Approximately 70% of genes, including *dlg-1*, receive an SL1 spliced leader that removes the 5′ portion of the primary transcript, thereby decoupling TSS position from mature 5′ UTR structure (11).

By analyzing single-copy integrated transcriptional reporters and CRISPR-generated deletion strains, we clarified the roles of the two HOT regions in the *dlg-1* locus during development. Specifically, a subdomain within the conserved region in the upstream HOT region is sufficient for wild-type transcriptional levels in all *dlg-1*-positive cells, and is required for normal development. The HOT region in the first intron possesses weak transcriptional capabilities that are restricted to epidermal cells, and are not essential for development. Together, our findings describe the physiological relevance of HOT regions containing TSS and/or enhancer sites and how they control general and selective gene expression in *C. elegans*. This study provides a physiological example of regulatory strategies used in *C. elegans* transcriptional regulation.

## Results and discussion

We focused our analysis of transcriptional regulation on the *dlg-1* locus, which is expressed in diverse tissues (6,7). The *dlg-1* promoter has been originally defined as a seven-kilobase (Kb) sequence that stretches from the CDS of the gene upstream on the minus DNA strand, *dpyd-1*, to the SL1 (believed to be also the TSS) of the *dlg-1* gene on the plus DNA strand (12) (Figure 1A and S1A). More recently, different promoters and TSS have been identified in the *dlg-1* locus (2,5,8). In analyzing the seven-Kb sequence upstream the *dlg-1* CDS, we identified a genomic region highly conserved among the *Caenorhabditis* species (Figure S1A). This region is bound by several transcription factors (TFs) suggestive of a high occupancy target (HOT) region (2,3) (Figure S1B). Large-scale studies aiming to identify HOT regions indeed picked up this genomic site as a HOT region and promoter of *dlg-1* (2,5,10). This HOT site also contains polymerase II peaks during embryogenesis (Figure S1C), confirming its role as a promoter (9). Controversially, *C. elegans* sequences that dictate correct expression levels and patterning are usually located within a few hundred base-pairs (bp) from a gene’s CDS (3,13– 15). The conserved region is positioned approximately four Kb from the CDS of *dlg-1* and is overlapping and bordered by chromatin marks typically found in enhancers. Specifically, it is enriched in acetylated lysine 27 (H3K27ac) and mono- and di-methylated lysine 4 of histone H3 (H3K4me1/2), and depleted from tri-methylated lysine 4 of histone H3 (H3K4me3) (Figure S1D) (5,16). Indeed, this genomic region has been predicted an enhancer site (5). Cohesins have been shown to organize the DNA in a way that active enhancers make contacts in the 3D space forming “fountains” detectable in Hi-C data (17,18). Although the presence of a fountain at this locus remains to be determined during embryogenesis, ARC-C data showed the putative HOT region interacts with other DNA sequences, including the first intron of *dlg-1* (1).

**Figure 1.**
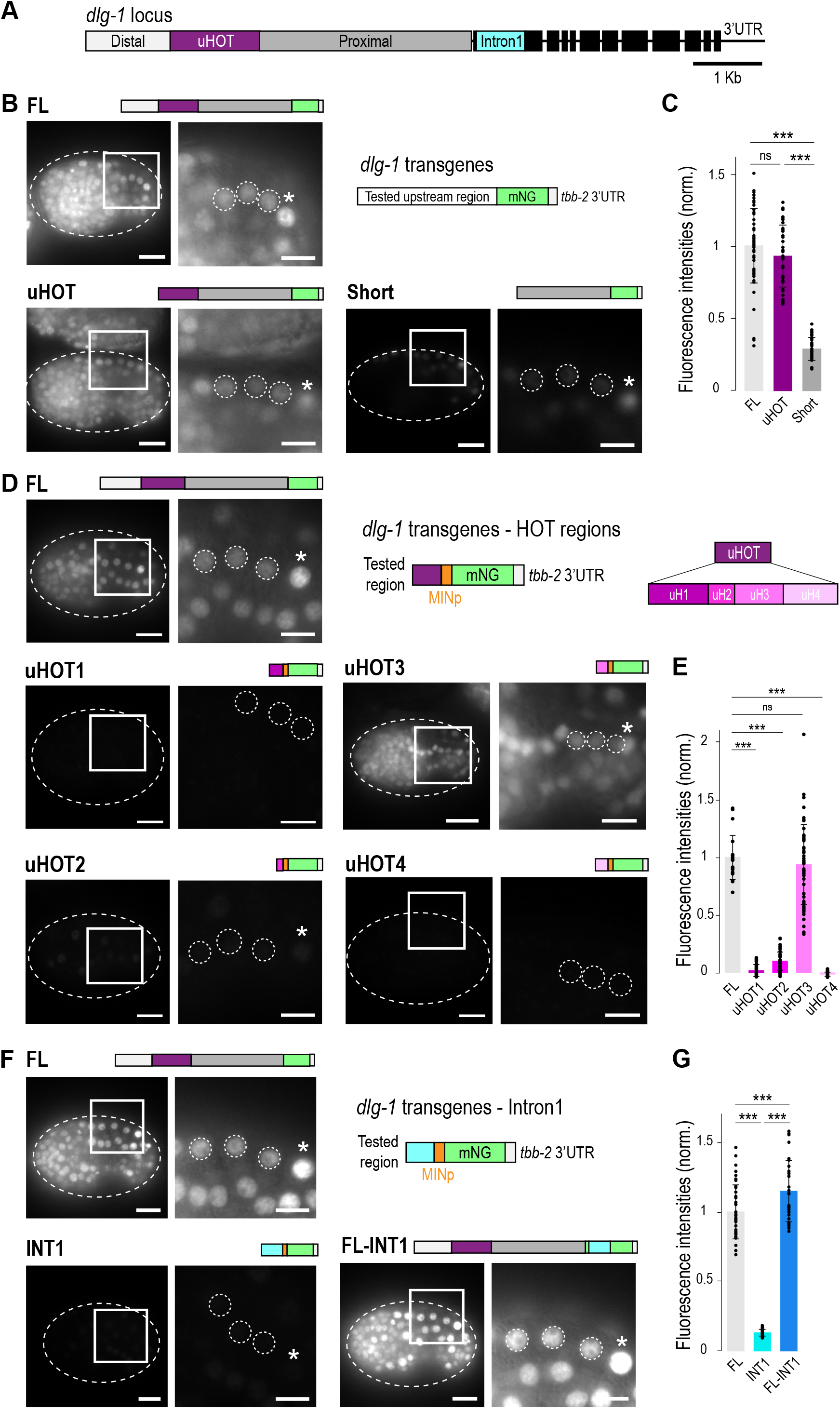
A small sequence within the upstream HOT region is the man driver of *dlg-1* transcription. **A**. Schematic representation of the *dlg-1* locus: black, exons (thick lines) and introns/UTR (thin lines); cyan, the first long intron; light grey, distal-most sequence of the upstream region to the CDS (Distal); dark purple, sequence in and around the upstream HOT region (uHOT); dark grey, proximal-most sequence of the upstream region (Proximal). Scale bar: 1 Kb. **B**. Upper-right side: schematic representation of the transgenic constructs used in this figure: H2B::mNG (mNG) coding sequence containing three synthetic introns (green) and a *tbb-2* 3’UTR (white) were under the control of upstream regions with different lengths (white). The fluorescent images are from live *C. elegans* embryos (outlined by dashed lines) at the comma stage and corresponding zoom-ins (squared boxes). The three seam cell nuclei used for quantitation are highlighted in the zoom-ins with dashed lines. A reference nucleus to identify the same measured nuclei in all the conditions is highlighted with an asterisk. The transgene expressed in the exemplifying embryos are stated above each image and accompanied with a small schematic. Scale bars: 10 µm (whole embryo) and 5 µm (zoom-in). **C**. Bar plot with error bars (standard deviation) normalized to the FL. Each dot represents the maximum fluorescent intensity of a seam cell nucleus. ns: not significant; ***: p < 0.01. For raw data, see Table S1. **D**,**F**. Upper-right side: schematic representations of the types of transgenic constructs used in these panels: H2B::mNG (mNG) coding sequence containing three synthetic introns (green) and a *tbb-2* 3’UTR (white) were under the control of portions of the upstream HOT region (purple, D) or the first intron (cyan, F) followed by a minimal *Δpes-10* promoter (MINp, orange). The fluorescent images are from live *C. elegans* embryos (outlined by dashed lines) at the comma stage and corresponding zoom-ins (squared boxes) of the three seam cell nuclei used for quantitation (smaller dashed lines depict nuclei boundaries). A reference cell to identify the same three seam cells measured in all the conditions in highlighted with an asterisk. The transgenes expressed in the embryos are stated above each image and accompanied with a small schematic. Scale bars: 10 µm (whole embryo) and 5 µm (zoom-in). **E**,**G**. Bar plot with error bars (standard deviation) normalized to the FL. Each dot represents the maximum fluorescent intensity of a seam cell nucleus (three nuclei per embryo). ns: not significant; ***: p < 0.01. For raw data, see Table S1.

We hypothesized the upstream HOT region is the main driver of *dlg-1* transcriptional regulation. To test this idea, we generated three single-copy, integrated transcriptional reporters carrying a histone H2B sequence fused to an mNeon-Green (mNG) fluorescent protein under the transcriptional control of the following *dlg-1* upstream regions: a full-length sequence (FL) and two deletion sequences spanning from the beginning (uHOT) or the end (Short) of the HOT region until the *dlg-1* CDS (Figure 1B,C). Live imaging and quantitation analyses showed minor and insignificant differences between the FL and the uHOT reporters regarding cell-type specificity (Figure 1B) and levels of expression (FL mean = 1.00, STDEV = 0.26; uHOT mean = 0.93, STDEV = 0.22; Figure 1C), and recapitulated that of the endogenous *dlg-1* gene (6,7). The Short reporter, containing nearly four Kb upstream the CDS and containing three other *dlg-1* promoters (5), showed a dramatic reduction in expression levels compared to FL and uHOT (Short mean = 0.29, STDEV = 0.08; Figure 1B,C). This reduction was observed in most of the cells where endogenous *dlg-1* is normally expressed (Figure 1B and S2). Only developing epidermal cells were still transcriptionally active, as shown previously with an extrachromosomal *dlg-1* transcriptional reporter with truncated upstream region from the CDS (19). smFISH experiments showed that the transcriptional control mediated by the upstream HOT region was specific to *dlg-1*, as the nearby *dpyd-1* locus was expressed sporadically (Figure S2). Together, these analyses demonstrate that the upstream HOT region is necessary to (i) increase majorly *dlg-1* expression in developing epidermal cells, and (ii) allow *dlg-1* expression in a broader range of cell types, such as neuronal and pharyngeal precursors. We conclude that the upstream HOT region constitutes a key regulatory sequence for correct expression levels and patterning of *dlg-1*.

To understand the sufficiency and determine which portion of the upstream HOT region was important for its function in gene expression strength and patterning, we selected four small regions within and surrounding the upstream conserved region (uHOT1 = 406 bp; uHOT2 = 185 bp; uHOT3 = 332 bp; uHOT4 = 374 bp) (Figure 1A,D,E). Each region was paired with the minimal promoter *Δpes-10* (MINp) (Figure 1D), commonly used to assess the functionality of candidate regulatory sequences in the context of transcription (20) and generated four transcriptional reporters. As previously described, we performed live imaging and fluorescence quantitation on these reporters to assess the levels and pattern of expression dictated by each sequence. Among the four synthetic promoters designed, only uHOT2 and uHOT3 were permissive for transcription (FL mean = 1.00, STDEV = 0.19; uHOT1 mean = 0.02, STDEV = 0.05; uHOT2 mean = 0.10, STDEV = 0.08; uHOT3 mean = 0.94, STDEV = 0.35; uHOT4 mean = −0.01, STDEV = 0.02; Figure 1D,E). The uHOT2 reporter showed scattered and variable expression that differed significantly in expression levels compared to the FL. The uHOT3 reporter showed patterning (Figure 1D) and expression levels (Figure 1E) comparable to the FL reporter. These data suggest that a small conserved sequence of 330 bp, which lies more than four Kb from the CDS, is a strong driver and main promoter of *dlg-1* transcription during embryonic morphogenesis.

We found a second region within the first long intron of *dlg-1* was also enriched for TF binding sites (3) (Figure S1B) and H3K4me1/2 chromatin modifications (13) (Figure S1D). Indeed the first intron has been reported to be a HOT region and contain promoter and enhancer sites (5). Long first introns have been shown to frequently contain regulatory regions used for transcription, such as enhancers (21–23). The enhancers located in introns can drive gene expression in patterns that are distinct from, yet complementary to, those found in the gene’s promoter (13). To test if the first intron contributed to *dlg-1* transcriptional control, we generated two single-copy integrated lines where the first intron was either paired with a minimal promoter *Δpes-10* (INT1) or inserted in the FL transgene within the H2B sequence, acting as an actual intron (FL-INT1) (Figure 1F). The reporter INT1 was created to test the transcriptional abilities of intron 1 alone, whereas the reporter FL-INT1 was engineered to determine if the first intron was changing the transcriptional capabilities of the upstream region (Figure 1F,G). We found that the fluorescence expression driven by the INT1 reporter was confined to developing epidermal cells (Figure 1F, S3) and minimal (13% of the FL; Mean FL = 1.00, STDEV FL = 0.19; Mean INT1 = 0.13, STDEV INT1 = 0.02; Figure 1G and S3), suggesting that this sequence plays a marginal role in *dlg-1* transcription. Interestingly, quantitation analyses on the FL-INT1 reporter showed a slight, but significant increase (+15%) in fluorescence intensity compared to the reference FL reporter in the tested cells (Mean INT1 = 1.15, STDEV INT1 = 0.22; Figure 1G). This result supports the idea of enhancers providing additive effect to gene transcription. We conclude that *dlg-1* intron 1 complements the main promoter, albeit weakly, in a subset of cells.

To clarify the impact of the two HOT regions during normal development, we generated two CRISPR lines derived from a strain with *dlg-1* tagged endogenously with a GFP sequence at the end of its CDS. The first line had the upstream HOT region removed (Δ-uHOT). Specifically, it spanned from bp 5,475,070 to 5,475,709 on chromosome X and removed HOT2 and HOT2 sequences. The second line had the first intron of *dlg-1* (spanning from bp 5,479,783-5,480,664 on chromosome X) substituted with a short synthetic intron of fifty-one bp (synt-INT1). We chose this synthetic intron as it is commonly employed in fluorescent reporter sequences to optimize their translation in *C. elegans*. Live imaging analyses showed a marked reduction to 46% (STDEV = 19) of DLG-1-GFP levels in Δ-uHOT compared to wild-type (wt) at the junction of embryonic epithelial cells - seam cells (Figure 2A,B and S4A). Furthermore, Δ-uHOT junctions presented gaps along their perimeter in these and other epithelial cells (Figure S4A,B). Single molecule fluorescent in situ hybridization (smFISH) experiments on fixed samples revealed these defects to be likely caused by a general reduction in *dlg-1* mRNA levels in all embryonic cells where *dlg-1* is normally expressed (Figure 2C and S1B). Such reductions in mRNA and protein levels determined severe although not fully penetrant developmental defects. Such defects include: reduced body (two days after L4 stage: wt mean = 1.19 mm; Δ-uHOT mean = 0.79 mm - Figure S4C,D) and brood sizes (wt mean = 164; Δ-uHOT mean = 28 - Figure S4E), and embryonic lethality or larval arrest (L1 stage) (survival to adult stage wt = 99.9%; Δ-uHOT = 3.9% - Figure S4F). To note, embryos managed to finish embryogenesis but die immediately before or shortly after hatching. This phenotype differs from both full knock-out of *dlg-1* where embryos survive until shortly after mid-embryogenesis (2-fold stage) (12) and its knock-down where animals develop normally, only at a slower pace (24). These results show that deletion of the upstream HOT region causes a reduction in *dlg-1* mRNA and consequently DLG-1-GFP protein levels in embryonic cells where *dlg-1* is normally expressed, leading to defects in junctional integrity and overall development. We conclude that the upstream HOT region is required for correct junction formation, epithelial integrity and normal animal development. Live imaging analyses on the synt-INT1 strain compared to wild-type (wt) did not reveal any differences at protein level (wt mean = 1.00, STDEV = 0.31; synt-INT1 mean = 1.03, STDEV = 0.30 - Figure 2D) or distribution of DLG-1-GFP in embryonic seam cells (Figure 2E), where intron 1 is permissive of transcription (Figure 1F and S3). Likewise, smFISH experiments did not show differences at level and distribution of *dlg-1* mRNA (Figure 2F). In line with these findings, synt-INT1 showed no evident phenotypes at any stage of development compared to wt. These data show that the replacement of the long intron 1 of *dlg-1* with a short synthetic intron does not alter mRNA or protein levels or distribution in embryonic seam cells. These results suggest that intron 1 is not required for *dlg-1* expression or function in cells where it however is permissive of transcription.

**Figure 2.**
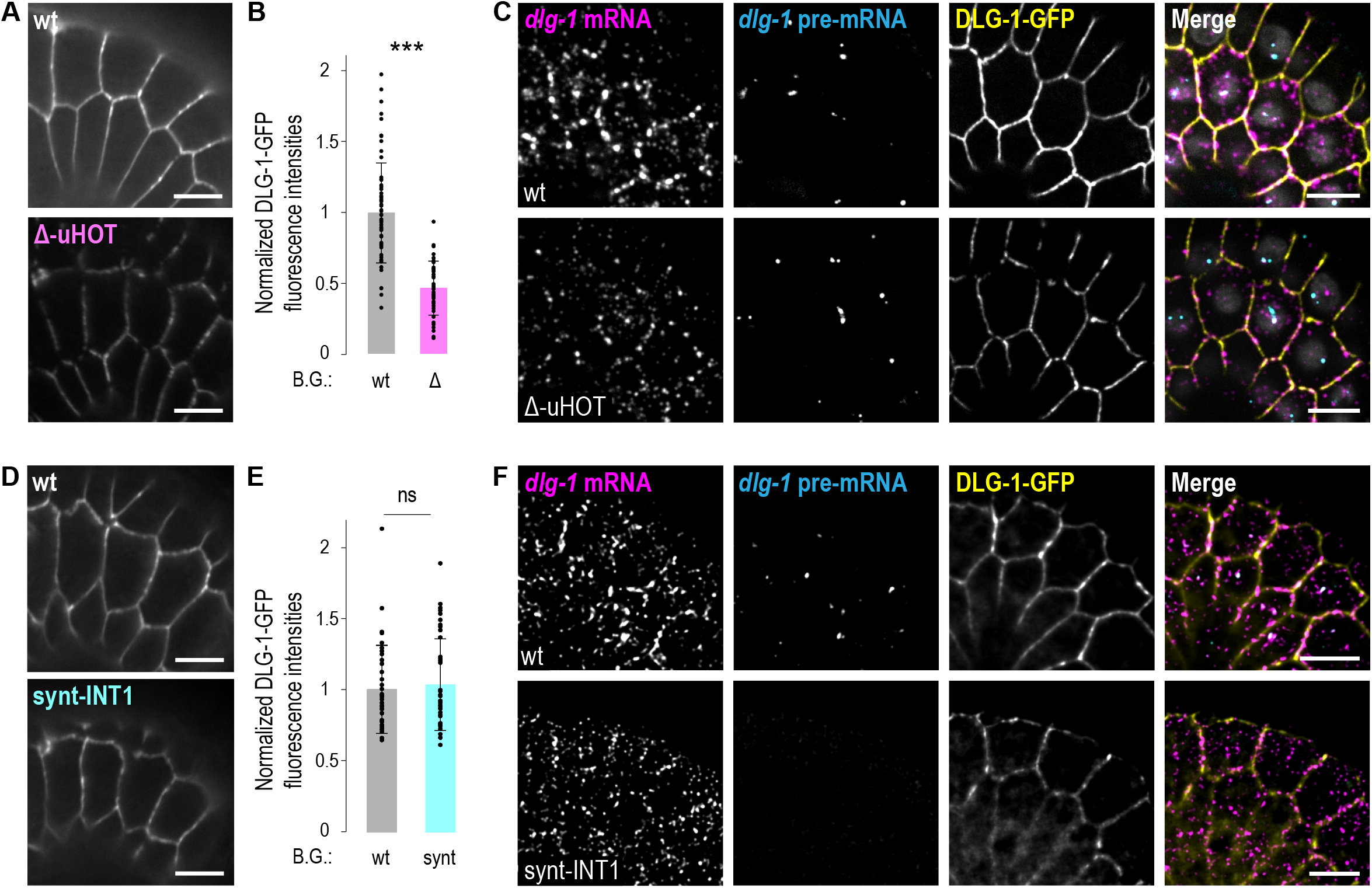
Deletion of the *dlg-1* upstream HOT region, but not intron 1, determines reduction at the mRNA and protein levels. **A,D**. Fluorescent images of zoom-ins on epidermal cells of live *C. elegans* embryos at the comma stage. The fluorescence comes from the GFP paired to endogenous DLG-1 in wild-type (wt, upper panel) or deletion strain for the upstream HOT region (Δ-uHOT). Scale bar: 5 µm. **B**,**E**. Bar plot with error bars (standard deviation) normalized to the wt. In (**B**) comparison of wild-type (wt, gray) to the deletion strain for the upstream HOT region (Δ, pink). In (**E**) comparison of wild-type (wt, gray) to the substitution strain for the first intron with a synthetic one (synt, light-blue). Each dot represents the maximum fluorescent intensity of a junction shared between two seam cells (five junctions per embryo – posterior-most ones). ns: not significant; ***: p < 0.01. For raw data, see Table S1. **C**,**F**. Fluorescent images of zoom-ins on epidermal cells of fixes samples of *C. elegans* embryos at the comma stage. Upper panels, wild-type strain (wt); lower panels, deletion strain for the upstream HOT region (Δ-uHOT, **C**) or substitution strain for the first intron with a synthetic one (synt-INT1, **F**). Panels from left to right: smFISH signal from *dlg-1* mRNA using probes complementary to exon region (magenta); smFISH signal from *dlg-1* pre-mRNA using probes complementary to intron regions (cyan); antibody stain signal from GFP fused to DLG-1 (yellow); and merges. Scale bar: 5 µm.

In summary, we described the roles of two key regulatory HOT regions in the *dlg-1* locus in modulating gene expression and development in the *C. elegans* embryo: a conserved portion of an upstream HOT region located over four Kb upstream of the CDS (HOT3), and another HOT region included in first long intron. HOT regions in *C. elegans* have been predominantly associated with promoters and were often considered non-functional or simply reflective of accessible chromatin. Our data provide clear functional evidence that the upstream HOT region can instead act as main driver of transcription and key for normal development. Specifically, we found that a 330 bp fragment within HOT3 is sufficient to drive strong and spatially patterned expression in embryonic cells, closely resembling the endogenous *dlg-1* expression profile. Deletion of the key regulatory sequence in the upstream HOT region determines a partial loss-of-function phenotype, mainly causing late embryonic lethality and L1 arrest. Nevertheless, such phenotypes are probably not the sole result of reduction in *dlg-1* mRNA, and consequently DLG-1 protein levels, as they are more severe than in other known mutants in which levels are affected, but the CDS of *dlg-1* is not (24). The upstream HOT region interacts with other genomic regions (1) and its depletion might affect gene expression of other genes beyond *dlg-1* itself. We also showed that the first intron acts additively with the upstream region to boost gene expression exclusively in developing epidermal cells in the context of transcriptional reporters. Despite that, exchanging the intron 1 with a synthetic one in the endogenous locus has not effect at the protein level, suggesting additional (co- or post-transcriptional) regulations to be in place to coordinate the translational output. The presence of shared TF binding both HOT/enhancer regions, HOT3 and the first intron of *dlg-1*, suggests that regulatory elements far upstream the CDS and intronic regions can function cooperatively. This discovery challenges previous reports that HOT regions are functionally isolated modules, and instead suggest a possible cooperative transcriptional regulation logic.

## Conclusions

Together, our findings illustrate how non-canonical regulatory elements (distal and intronic HOT regions) are integrated to fine-tune gene expression in a cell-type-specific manner during development. These results help to clarify the functional potential of HOT regions in *C. elegans* and expand our understanding of how promoter and enhancers contribute to precise spatiotemporal control of gene expression in compact genomes like the one of *C. elegans*.

## Methods

### Nematode culture

*C. elegans* strains were maintained as previously described (25) at 20°C. Embryos were imaged at the comma stage. For a detailed list of transgenic lines used in this study, see Table S1.

### Generation of transgenic lines

Transgenes carrying the different promoters analyzed in this study (amplified from N2 lysate) were fused to an H2B sequence (amplified from N2 lysate) paired to an mNG sequence through a flexible linker (amplified from LP598 lysate), and a *tbb-2* 3’UTR (amplified from N2 lysate). These sequences were assembled (NEBuilder® HiFi DNA Assembly Cloning Kit, New England BioLabs, cat#E5520) in the pCFJ150 vector and inserted into the *ttTi5605* locus on chromosome II of the mosSCI strain EG6699 (26). NEB® 10-beta Competent *E. coli* (High Efficiency) (NEB, cat#C3019) competent cells were used for transformation. Full plasmid sequencing from Microsynth was used to confirm the sequences were correct. Plasmids were purified with MIDI preps (NucleoBond Xtra Midi Plus (Macherey - Nagel, cat#740412.10) and single-copy integrated lines were generated following the mosSCI protocol (26). For a detailed list of plasmids used in this study, see Table S1.

Upstream HOT site deletion and synthetic intron 1 replacement lines were obtained using the CRISPR/Cas9 technique, following a previously described protocol (27). For a complete list of reagents (crRNA, repair oligos, and genotyping/sequencing primers), see Table S1.

### smFISH

smFISH experiments were performed as previously described (7,24,28). For a full list of smFISH probes, see Table S1.

A widefield microscope FEI “MORE” with total internal reflection fluorescence (TIRF), equipped with a Hamamatsu ORCA flash 4.0 cooled sCMOS camera, and a Live Acquisition 2.5 software were used for capturing smFISH images. Images were deconvolved with the Huygens software and processed in OMERO (https://www.openmicroscopy.org/omero/). All figures were assembled in Adobe Illustrator (https://www.adobe.com/).

### Live imaging: microscopy, quantitation, and analysis

Live imaging was performed on embryos at the comma stage. Embryos were washed from plates with 1 ml of water, collected into Eppendorf tubes, and spun-down (“short”) for 5 seconds in a table centrifuge. After removing the supernatant, embryos were resuspended in M9 buffer and transferred onto poly-L-lysine-coated slides (Thermo Scientific, cat#ER-308B-CE24). A coverslip was placed on the sample and embryos were imaged immediately.

A widefield ZEISS Axio Imager M2 equipped with a ZEISS Colibri LED light source, a Hamamatsu ORCA flash 4.0 camera, and ZEN 2.6 software (blue edition) was used for capturing images for live imaging of *C. elegans* embryos.

Fluorescence signal quantitation of nuclear (from H2B-mNG) and junctional (from DLG-1-GFP) signals in the different transgenic lines was performed using Fiji, where the maximum intensity was quantified.

Statistical analyses were performed using R software (R Core Team, 2021; https://www.R-project.org/). The ggplot2 package (Wickham, 2016; https://ggplot2.tidyverse.org) was used to generate bar plots with error bars (standard deviation) and dot plots with box plots. Each dot represents the normalized value to the corresponding seam cell nucleus or junction. Statistical differences were determined using a *t*-test. The clustered column chart was created with Microsoft Excel.

## Declarations

### Ethics approval and consent to participate

Not applicable.

### Consent for publication

Not applicable.

### Availability of data and materials

All data are available in the described sources in the Methods.

### Authors’ contribution

Conception of the work (CT and SEM); design of the work (CT); acquisition and analysis (CT, FE and PB); interpretation of data (CT); drafted the work (CT); revised the work (SEM). All authors have read and approved the current version of this manuscript and have agreed to its submission for publication.

## Acknowledgments

We thank current and previous lab members of the Mango group for scientific discussions, the Imaging Core Facility (IMCF) of the Biozentrum for technical support, and WormBase. A particular thank to Dr. Camila Pulido for reading and giving comments to the manuscript. Some strains were provided by the CGC, which is funded by the NIH Office of Research Infrastructure Programs (P40 OD010440).

