## Supplemental Data for "Differential functions of upstream and intronic HOT regions shape *dlg-1* transcription and *C. elegans* development"

### Supplemental information

#### Supplemental figure legends

**Figure S1. Enrichments of TFs, PolII, and enhancer chromatin modification from ChIP data in *dlg-1* upstream HOT region and intron 1. A-D.** Snapshots from JBrowse of part of the *dlg-1* locus: from the dpyd-1 TSS to the beginning of the second exon of *dlg-1*. Tops and bottoms of the different panels show curated genes and schematics used to generate the transgenic lines in this study, respectively. In (**A**): conservation. In (**B**): ChIP experiments on embryos of TFs listed above the peaks (1). In (**C**): ChIP experiments on embryos of PolII at the listed developmental stages (2). In (**D**): ChIP-sequencing (ChIP-seq) experiments on embryos of four chromatin marks (H3K27Ac, H3K4me1/2/3) (1). Magenta squares focus on the upstream HOT region and surrounding sequences, and cyan squares on the first intron of *dlg-1* containing the second HOT site.

**Figure S2. The *dpyd-1* gene has a different expression pattern than *dlg-1*.** Fluorescence images of a *C. elegans* embryo at the comma stage. Upper panels, section of the surface of the embryo; lower panels, section of the middle part of the same embryo. Panels from left to right: smFISH signal from *dlg-1* mRNA using probes complementary to exon region (magenta); smFISH signal from *dpyd-1* pre-mRNA using probes complementary to intron regions (cyan); antibody stain signal from GFP fused to DLG-1 (yellow); and merges. Scale bar: 10 µm.

**Figure S3. The *dlg-1* intron 1 sequence followed by a minimal promoter drives expression in embryonic epidermal cells.** Fluorescent images of live *C. elegans* embryos (outlined by dashed lines) at the comma stage. The left and right panels show the same embryos but at different exposure intensities (1x left, 5x right) to better visualize the fluorescence-positive nuclei in INT1. The transgene expressed in the embryos are stated above each image (FL and INT1) and accompanied with a small schematic. Scale bar: 10 µm.

**Figure S4. Deletion of the *dlg-1* upstream HOT region determines problems at junctional levels and distribution, mRNA abundance, and developmental defects. A.** Clustered column chart of normalized intensity data from endogenously-tagged DLG-1 with GFP along the junction (perimeter) of one example seam cell from wild-type (wt, gray) and deletion strain for the upstream HOT region (Δ-uHOT, pink). Each column represents the maximum intensity at the stated location along the perimeter of the seam cells. Two purple arrows highlight the presence of gaps along the junction of the Δ-uHOT seam cell. **B.** Fluorescent images of zoomins on pharyngeal cells of fixes samples of the same *C. elegans* embryos in Figure 2C. Upper panels, wild-type strain (wt); lower panels, deletion strain for the upstream HOT region (Δ-uHOT). Panels from left to right: smFISH signal from *dlg-1* mRNA using probes complementary to exon region (magenta); smFISH signal from *dlg-1* pre-mRNA using probes complementary to intron regions (cyan); antibody stain signal from GFP fused to DLG-1 (yellow); and merges. Scale bar: 5 µm. **C.** Bright field images of free living *C. elegans* wt and Δ-uHOT adult worms (2-days after L4 stage) on NMG plates seeded with OP50 bacterial strain. Scale bar: 100 µm. **D-F.** Dot plots with box plots. Each dot represents: (**D**), the body length in µm of one wt (wt, gray) or Δ-uHOT (Δ, pink) adult animal (2 days after the L4 stage); (**E**), the brood size from the animals whose length was quantified in (**D**); (**F**), the survival rate to adulthood of the progeny quantified in (**E**). For raw data, see Table S1. Significance of statistical analyses (*t*-test, two tails): ***: $p < 0.01$.

#### Supplemental table legends

**Table S1. Quantitation and reagent lists.** Raw data regarding Figures 1C,E,G, 2B,E, and S4A,D-F. List of reagents used for the generation of CRISPR lines, *C. elegans* strains (3–5), plasmids, and smFISH primary probes used in this study. Available with DOI: 10.5281/zenodo.16762430.
