## Supplementary figures and images for "Differential functions of upstream and intronic HOT regions shape *dlg-1* transcription and *C. elegans* development"

### Figure S1

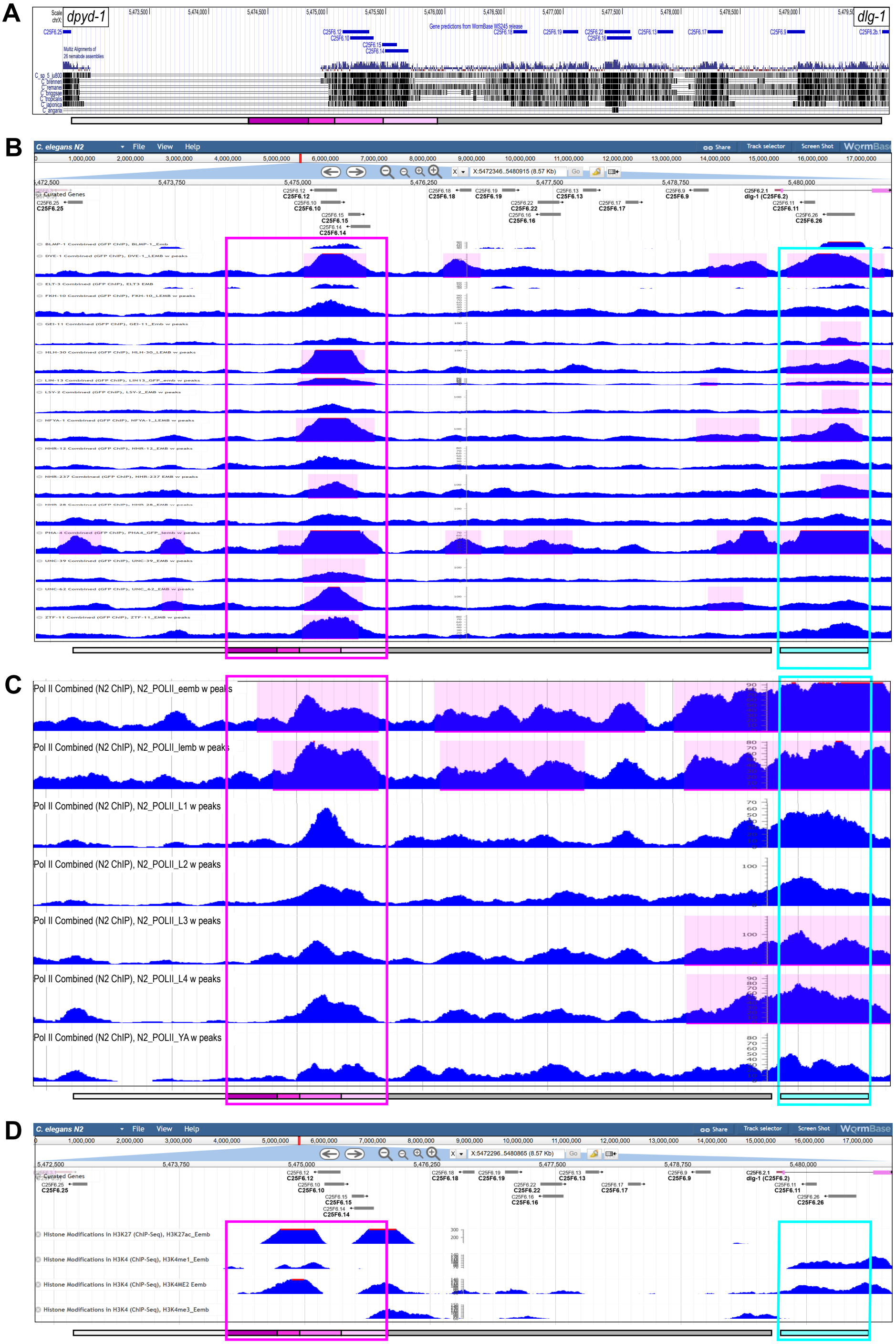


Fig.S1, Tocchini *et al.*

### Figure S2

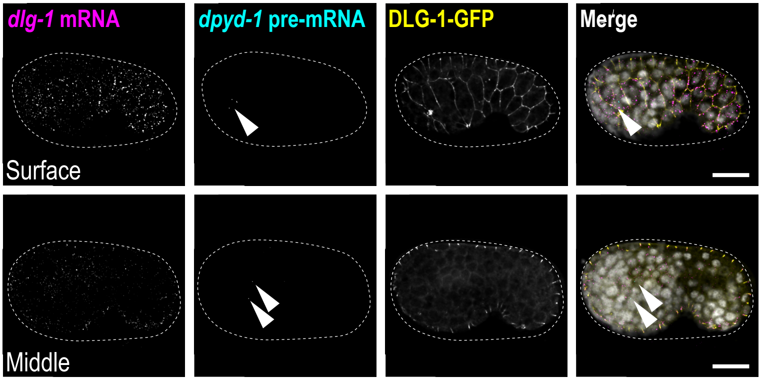


Fig.S2, Tocchini *et al.*

### Figure S3

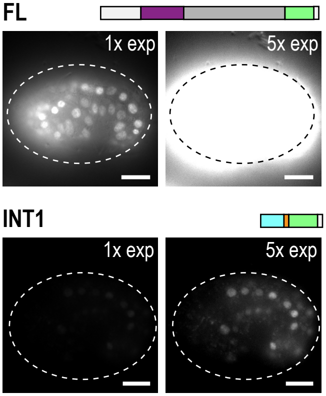


Fig.S3, Tocchini *et al.*

### Figure S4

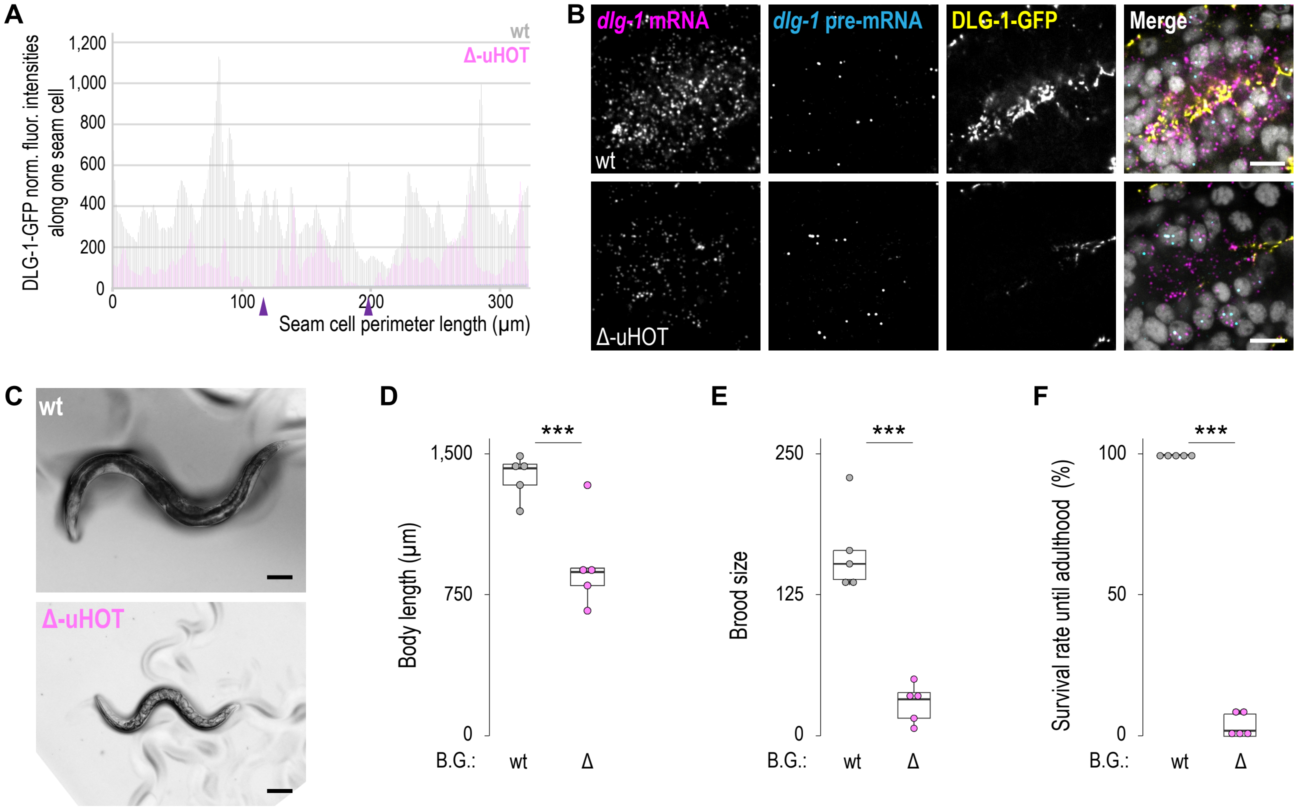


Figure S4_Tocchini *et al.*
